# Ex Vivo Herpes Simplex Virus Reactivation Involves a DLK-Dependent Wave of Lytic Gene Expression that is Independent of Histone Demethylase Activity and Viral Genome Synthesis

**DOI:** 10.1101/2022.02.25.481951

**Authors:** Abigail L. Whitford, Corinne A. Clinton, E.B. Lane Kennedy, Sara A. Dochnal, Jon B. Suzich, Anna R. Cliffe

## Abstract

Herpes Simplex virus-1 (HSV-1) maintains a lifelong latent infection in neurons and periodically reactivates, resulting in the production of infectious virus. The exact cellular pathways that induce reactivation are not understood. In primary neuronal models of HSV latency, the cellular protein Dual Leucine Zipper kinase (DLK) has been found to initiate a wave of viral gene expression known as Phase I. Phase I occurs independently of both viral DNA replication and the activities of histone demethylase enzymes required to remove repressive heterochromatin modifications associated with the viral genome. Here we investigated whether Phase-I like gene expression occurs in ganglia reactivated from infected mice. Using the combined trigger of explant-induced axotomy and inhibition of PI3K signaling, we found that HSV lytic gene expression was induced rapidly from both sensory and sympathetic neurons. *Ex vivo* reactivation involved a wave of viral late gene expression that occurred independently of viral genome synthesis and histone demethylase activity, and preceded the detection of infectious virus. Importantly, we found that DLK was required for the initial induction of lytic gene expression. These data confirm the essential role of DLK in inducing HSV-1 gene expression from the heterochromatin associated genome and further demonstrate that HSV-1 gene expression during reactivation occurs via mechanisms that are distinct from lytic replication.

## Introduction

The ubiquitous human pathogen Herpes Simples Virus persists for life in the form of a latent infection in neurons. In response to a variety of different stimuli, the virus can reactivate from the latent state, resulting in the release of infectious virus and subsequent replication in the surrounding tissue. Clinically, reactivation of the virus can manifest as a variety of disease states including lesions at the body surface, keratitis, and encephalitis. In addition, there is growing evidence of a link between HSV infection and the development of late onset Alzheimer’s disease, particularly in individuals with the ApoE4 variant (1-9). There are potentially different stimuli that can induce HSV to reactivate from latency. These stimuli may converge on single cellular pathways or operate via distinct mechanisms to induce reactivation (10). Identifying the cellular pathways important in reactivation is required to understand how viral gene expression initiates from the latent genome and to ultimately develop therapeutics to prevent reactivation occurring.

During a latent infection of neurons, the HSV genome is assembled into repressive heterochromatin. This has been characterized by the enrichment of post-translational modifications on histone H3; namely di- and tri-methyl lysine 9 (H3K9me2/3) and tri-methyl lysine 27 (H3K27me3) on lytic promoters (11-17). By assembling into heterochromatin, viral lytic transcripts are maintained in a silent state. In addition, both host and viral miRNAs target lytic mRNAs (18-22). Therefore, the action of transcriptional and translational silencing results in limited synthesis of viral lytic proteins. This lack of lytic proteins suggests that HSV is reliant upon host signaling to initiate gene expression and reactivation. To understand the mechanism of HSV reactivation, it is important to determine how activation of host cell pathways ultimately converge on the repressed viral genome to induce lytic gene expression.

There is evidence that the initial induction of HSV-1 lytic gene expression following reactivation occurs in a manner that is distinct from the mechanisms of viral gene expression during lytic replication. HSV lytic genes can be divided into groups characterized by their requirements for viral protein synthesis and viral DNA replication during lytic replication (23). Immediate early (IE) genes are expressed independently viral protein synthesis during lytic infection and instead require the viral tegument protein VP16 for maximal expression. Early (E) genes are expressed following the production of IE proteins. Certain IE proteins (including ICP4, ICP0 and ICP27) stimulate viral E protein synthesis. Late (L) genes require viral DNA synthesis and are subdivided into genes that are expressed at low levels even prior to DNA replication but increased with genome synthesis (leaky L) and those that are fully dependent on viral DNA replication (true L). The dependence on DNA replication for L gene expression is not fully understood but likely involves a shift in genome accessibility and increased binding of host transcriptional machinery (RNA Pol II, TBP, and TAF1) (24). In contrast to this regulated cascade, in models of HSV reactivation, an initial burst of lytic gene expression, named Phase I of reactivation, has been observed where inhibition of protein synthesis prior to the accumulation of IE transcripts does not prevent E gene expression (25). In addition, L gene expression is unaffected by inhibition of viral DNA replication (25). Together, these data indicate that initial expression of lytic transcripts during the early stages of reactivation does not resemble the early stages of *de novo* infection in non-neuronal cells.

Phase I of HSV reactivation has largely been identified in primary neuronal models of HSV latency (26). In these experimental models, a variety of stimuli have been found to induce HSV to reactivate from a latent infection, including loss of neurotrophic factor support (25, 27-29), increased neuronal excitation (30), modulation of DNA damage/repair (31) and exposure to corticosteroids (27). Precisely how activation of these pathways permits expression of the viral gene transcripts for reactivation to occur is not fully understood. Previously, we have identified a role for the cell stress protein, dual leucine zipper kinase (DLK), in inducing Phase I of HSV reactivation (27). DLK is activated by the loss of neurotrophic factor support, which can be mimicked by inhibition of Phosphoinositide 3 (PI3)-kinase activity (27), and during heightened neuronal excitation and interleukin-1 (IL-1) treatment (30). DLK is a master regulator of axonal responses to stress and can mediate a variety of responses including Wallerian degeneration, axon regeneration, apoptosis, and axon pruning (32). Upon activation, DLK is known to re-direct the cell stress protein, c-Jun N-terminal kinase (JNK), from its physiological role in neurons, maintaining synaptic arborization, to its cell stress function (33). Accordingly, we and others have identified a role for JNK in reactivation of HSV from latency (27, 30, 31). JNK is also important in reactivation of the related Alpha herpesvirus, Varicella-Zoster virus, from a latent infection (34) highlighting its potential central role in reactivation of human Alpha herpesviruses.

A role for DLK and JNK in HSV reactivation has been mostly widely studied on primary neuronal models of latency in murine sympathetic neurons (27, 30). In these neurons, latency is established in the presence of the HSV DNA replication inhibitor, acyclovir. After the removal of acyclovir, reactivation can be induced by PI3-kinase inhibition or forskolin (25, 27, 30, 35, 36). In these systems, full reactivation occurs around 48h post-stimulus, which requires the activities of histone demethylase enzymes, indicating that reactivation requires removal of repressive heterochromatin (27, 30). However, the DLK-dependent Phase I peaks around 18h post-stimuli and (27, 30, 37), importantly, Phase I occurs independently of lysine 9 and lysine 27 histone demethylase activity (27, 30). Instead, JNK activation induces histone phosphorylation on H3Serine10 (H3S10) on histones that maintain the H3K9me3 modification (27); this is known as a histone methyl/phospho switch and presumably permits transcription by overriding the repressive H3K9me3 modification. However, although demonstrated *in vitro*, the possibility of a DLK/JNK-dependent wave of gene expression, characteristic of Phase I, has not been explored following *in vivo* infections.

A robust mode of HSV reactivation from infected mice is explanation of the sensory trigeminal ganglia (TG). The action of severing the axon (axotomy), is thought to be the trigger that induces HSV to reactivate. In this model, whether a Phase I-like wave of gene expression occurs has not been fully explored. Data from a previous study suggests that a H3 lysine 9 demethylase functions to promote lytic gene expression at an early time-point (6 hours post-explant) (38). This indicates that wave of gene expression that is independent of histone demethylase activity occurs at an earlier time-point, or that explant induced reactivation does not involve a Phase-I like wave of viral gene expression. There are multiple differences between the *in vivo* experiments and *in vitro* latency models, including the presence of host immune system *in vivo*. In addition, explant-induced reactivation has been investigated mostly in sensory neurons whereas models for *in vitro* infection often use sympathetic neurons; both neuronal types are targets of HSV latent infection in humans (2, 39-41). *In vitro* models also use acyclovir to promote the establishment of latency. Finally, potential differences resulting from viral strains used cannot be ruled out. In experiments investigating the role for histone demethylases *ex vivo*, HSV strain F was used (16, 38), whereas the in vitro models have been performed with the KOS and Patton strains (25, 27).

To determine whether previous observations of a DLK/JNK triggered Phase I reactivation *in vitro* were recapitulated e*x vivo*, we dissected trigeminal ganglia from latently infected mice and determined whether we observed lytic gene expression in response to PI3-kinase inhibition. We found that after only five hours post excision, in treated ganglia, there was robust expression of IE, E, and L viral genes and this gene expression was independent of viral DNA replication. Supporting previous findings, we also found that this initial burst of lytic gene expression was not dependent upon LSD1 (H3K9-demethylase) nor JMJD3 and UTX (H3K27-demethylases). Therefore, Phase I of reactivation was observed *ex vivo* and was not reliant upon the removal of repressive heterochromatic marks. We found that this Phase 1 was dependent upon DLK, thus indicating that neuronal stress pathways can trigger bi-phasic reactivation of HSV-1 in ganglia *ex vivo*.

## Materials Methods

### Cells and Viruses

HSV-1 stocks of KOS were grown and titrated on Vero cells obtained from the American Type Culture Collection (Manassas, VA) as described previously (11). Cells were maintained in Dulbecco’s Modified Eagle’s Medium (Gibco) supplemented with 10% FetalPlex (Gemini Bio-Products) and 2 mM L-Glutamine. KOS was kindly provided by Dr. David Knipe, Harvard Medical School.

### Reagents

Compounds used in the study are as follows: Acycloguanosine (acyclovir; ACV), GSK-J4 and GNE-3511(Millipore Sigma) SP600125 (Thermo Fisher Scientific), OG-L002 (Tocris); Nerve Growth Factor 2.5S (Alomone Labs).

### Mouse Infections

Six-week-old male and female CD-1 mice (Charles River Laboratories) were anesthetized by intraperitoneal injection of ketamine hydrochloride (80mg/kg) and xylazine hydrochloride (10mg/kg) and inoculated with 1.5 × 10^6^ PFU/eye of virus (in a 5μl volume) onto scarified corneas, as described previously (11). Mice were housed in accordance with institutional and National Institutes of Health guidelines on the care and use of animals in research, and all procedures were approved by the Institutional Animal Care and Use Committee of the University of Virginia. Criteria used for clinical scoring based on the formation of lesions, neurological and eye symptoms is shown in table 1 and based on a previously establishing scoring scale (42). Mice were randomly assigned to groups and all experiments included biological repetitions from independent litters.

**Table 1.** Clinical scoring scale for HSV infected mice.

| Score | Description of score by type |  |  |
| --- | --- | --- | --- |
|  | Lesion | Eye | Neurological |
| 0 | Lesion free | Symptom free | Symptom free |
| 1 | Small area of broken skin <0.5cm | Pus around edges | Hunched posture, normal movement |
| 2 | Area of broken skin 0.5-1 cm | Pus and squint | Hunched posture, slow movement |
| 3 | Bleeding, scabbing or pustules | Closed | Hunched posture, labored breathing and/or hind leg paralysis |
| 4 | Broken skin >cm with multiple pustules or scabbing | Scab formation | Hunched, ruffled fur, little to no movement |
| 5 | Sever scabbing or bleeding with pustules | Sever scabbing | Moribund or dead |

### Explant Induced Reactivation

Trigeminal ganglia (TG) and superior cervical ganglia (SCG) were removed at least 28 days post-infection. The ganglia were maintained intact and immediately placed in reactivation media containing DMEM/F12 (Gibco) supplemented with 10% Fetal Bovine Serum and Mouse NGF 2.5S (50 ng/mL). Compounds were added at the following concentrations; LY294002 40μM, GNE-3511 8μM, ACV 100μM, GSK-J4 10μM, OG-L002 50μM. For treatments with GNE-5311, ACV, GSK-J4 and OG-L002, ganglia were treated with the compounds 3-60 minutes prior to the addition of LY294002. Ganglia were placed on a shaking platform at 50rpm at 37°C.

### Quantification of Viral Transcripts and Genome Copy Number

At the required time point, the reactivation media was removed and ganglia snap-frozen in liquid nitrogen. Lysis was carried out by addition of the excised ganglia to BeadBug™ homogenization microtubules then homogenized for 60 seconds using the BeadBug™ microtube homogenizer. DNA and RNA were isolated from the homogenized mixture using the *Quick* DNA/RNA miniprep kit. Following RNA isolation TURBO DNA-free Kit (Invitrogen) was used to remove any contaminant DNA. mRNA was reverse transcribed into cDNA using MaximaRT (ThermoFisher) using random hexamers for first strand synthesis. Equal amounts of mRNA were used for each reverse transcriptase experiment (20-30 ng/reaction). Power SYBR Green PCR Master Mix was used for qPCR (Applied Biosystems). Standard Curves were used to calculate the relative mRNA or DNA copy number per genome and were standardized to cellular GAPDH or 18s. All RNA samples were run in triplicate and all DNA samples were run in duplicate on an Applied Biosystems™ QuantStudio™ 6 Flex Real-Time PCR System. The primers that were used are described in Table 2.

**Table 2.**
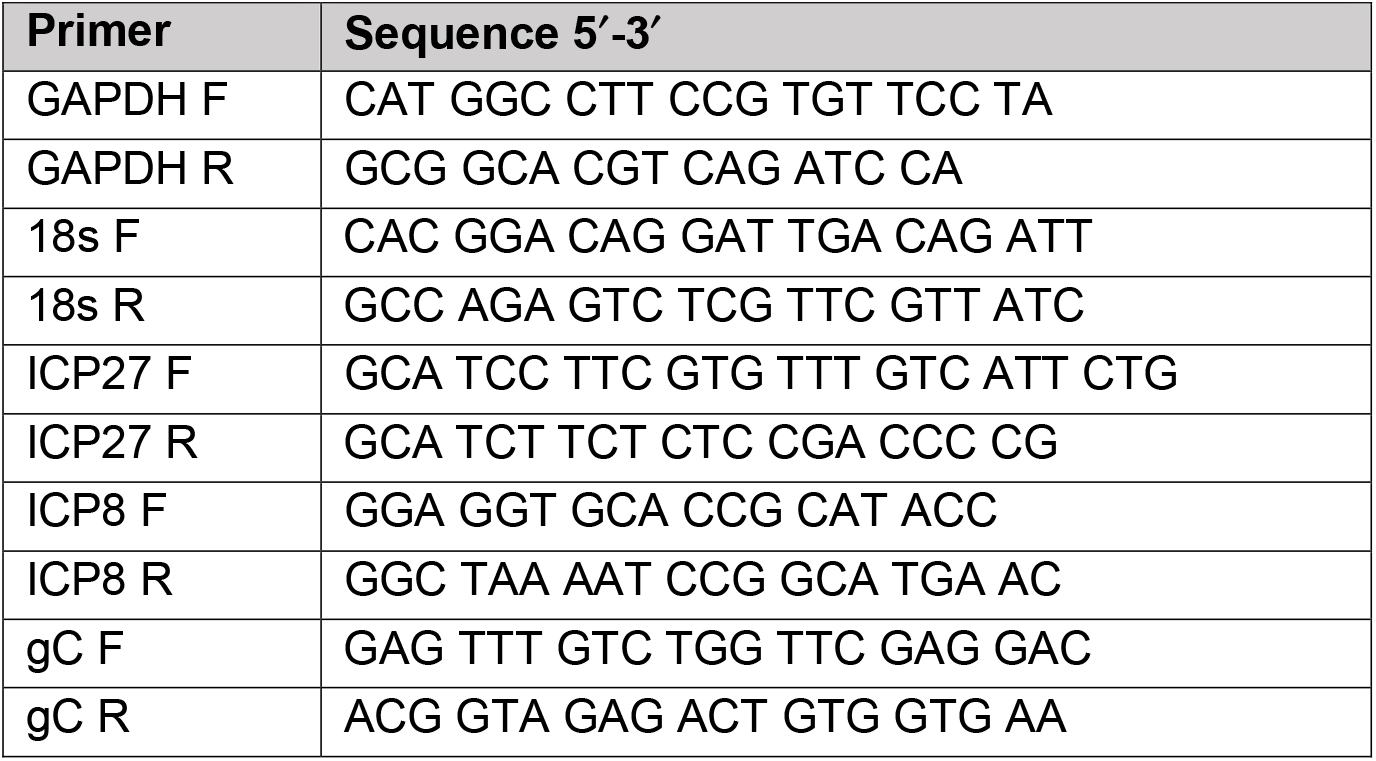
Oligonucleotides used in this study

### Preformed virus titration

Sterile milk was added to media containing reactivated ganglia and sample snap frozen. Ganglia were thawed at 37°C then homogenized using a BeadBug microtube homogenizer. The homogenized ganglia were then sonicated at 25% for 30 seconds three times on ice in a Fisherbrand™ Model 120 Sonic Dismembrator. After sonication, ganglia homogenates were frozen on dry ice and thawed at 37°C. Homogenates from all time points were simultaneously titrated on Vero cells.

## Results

### Explant combined with PI3-kinase inhibition triggers robust HSV-1 lytic gene expression

We first set out to determine whether a wave of lytic gene expression occurs following reactivation of explanted ganglia, similar to what has been observed in primary neuronal cultures. Female mice were infected via the ocular route and reactivation studies were carried out at least 28 days post-infection. Previously, studies from our lab and others have shown that addition of a reactivation stimulus to sympathetic neurons infected *in vitro* yields induction of lytic viral gene expression around 15-20 hours post-stimulus (25, 27). In addition, a previous study examining explant induced reactivation combined with deprivation of nerve growth factor (NGF) also resulted in lytic gene induction around 12-15 hours post-explant (43). Therefore, we initially examined the effect of PI3-K inhibition (using LY294002; 40 μM) at 20 hours post-reactivation to determine whether loss of the PI3K/AKT branch of the NGF-signaling pathway also promotes reactivation *ex vivo*. Acyclovir (ACV; 100 μM) was also introduced alongside LY294002 because late gene expression during the initial activation of viral gene expression in *in vitro* models has been reported to proceed independently of DNA replication (25).

In ganglia that were explanted and maintained in NGF to provide continued neurotrophin support, very little induction of lytic gene expression was observed at 20h post-explant, especially for representative IE and E transcripts. A slight increase in *gC* mRNA was observed, although this increase was not statistically significant (P=0.1255, Mann Whitney U Test). However, the addition of the PI3-kinase inhibitor (LY294002) increased viral gene expression by 100-1000-fold. RT-qPCRs were carried out using primers against IE (ICP27, Fig.1 B), E (ICP8, Fig. 1C) and L (gC, Fig. 1D) genes. All three genes were significantly induced at 20h post excision. The addition of ACV to inhibit viral DNA replication did not inhibit the induction of IE or E gene expression, as expected. However, expression of the gC mRNA was significantly reduced in the presence of ACV, indicating that its maximal expression was dependent on viral genome synthesis. However, gC mRNA levels with ACV were still significantly increased compared to the latent samples. At 20h post-reactivation, an increase in viral genomes was also observed (Fig. 1A). Together, these data show that at 20h post-reactivation viral gene expression does not resemble the Phase I observed in primary neuronal cultures, as genome synthesis has occurred, and full late gene expression is dependent on viral DNA replication. However, this did not rule out the possibility of a Phase I-like wave of gene expression occurring prior to 20h, especially as *gC* mRNA was still induced compared to the 0h (latent) samples, indicating that there may still be some *gC* expression occurred even from non-replicated viral genomes.

**Figure 1.**
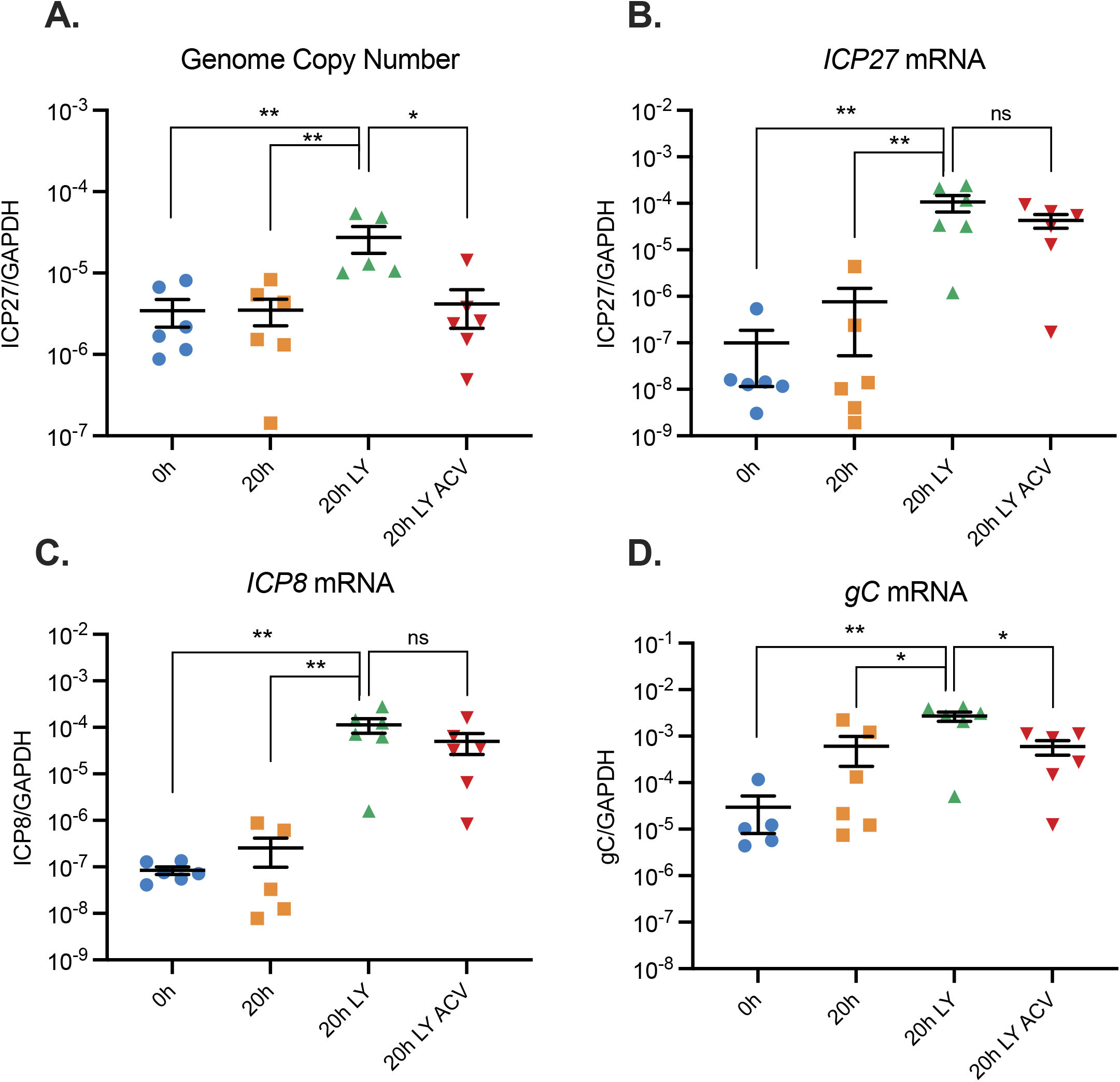
Explant combined with PI3-kinase inhibition triggers robust HSV-1 lytic gene expression. Mice were infected via corneal scarification and at least 28 days post infection trigeminal ganglia (TG) were excised. Ganglia were either snap frozen at 0 hours, or reactivated for 20 hours in media alone, with LY294002 (40 μM) or LY294002 with acyclovir (ACV; 100μM). (A) Viral genome copy number was quantified by qPCR. (B) Viral gene expression was quantified by RT-qPCR for immediate early (*ICP27*), (C) early (*ICP8*) and (D) and late (*gC*) genes. Transcript copy number was normalized to cellular control (*GAPDH*). n=6 biological replicates. Mann-Whitney U (A-D). *p<0.05, **p<0.01. Individual biological replicates along with the means and SEMs are represented.

### PI3K inhibition/explant induces rapid late gene expression in the absence of detectible genome synthesis in trigeminal ganglia

To determine whether viral lytic gene expression occurred prior to 20h post-stimulus in explanted ganglia with features characteristic of Phase I gene expression, we decided to undertake both male and female mice and determine changes in lytic gene expression following PI3-kinase inhibition of explanted ganglia. Because we infected both male and female mice, we assessed if sex impacted clinical manifestations of infection, mice were monitored throughout the period of infection. Both male and female mice had minimal mortality with only 92 and 88 percent mortality respectively. The mortality rates between sexes were not found to be significant by Kaplan-Meier survival analysis (Fig. 2A). To further analyze clinical symptoms, mice were evaluated by scoring lesions, neurological symptoms and eye health based on a previously described scoring metric (42). Infection of female mouse resulted in more severe clinical manifestations at days 7 and 9 post-infection (Fig. 2B). Despite these differences, there was no significant differences in the percentage of weight loss following infection between male and female mice (Fig. 2C). Based on these criteria, we concluded there were only minor sex-dependent differences upon HSV infection.

**Figure 2:**
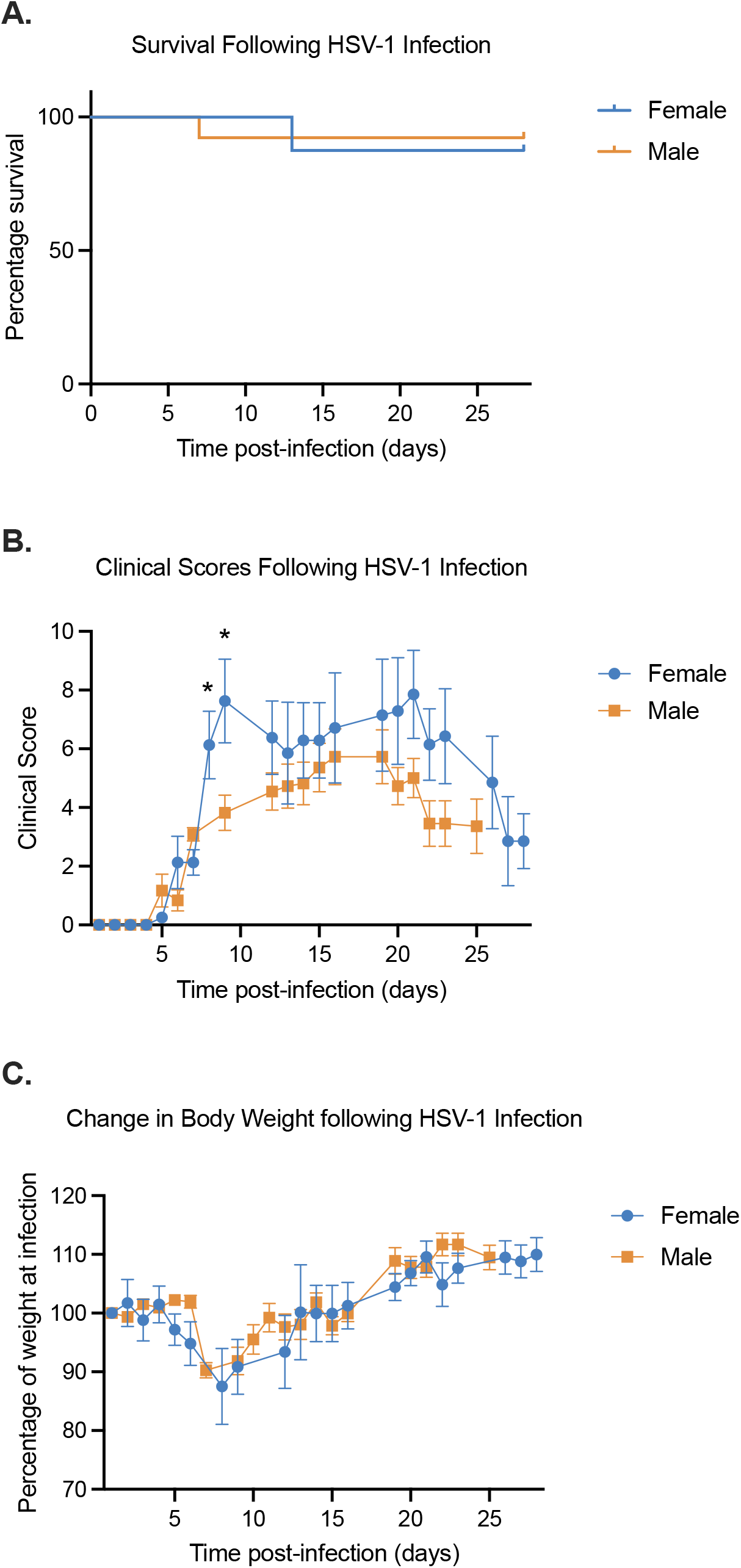
Sex dependent phenotypes during HSV-1 *in vivo* infection. Mice were infected via corneal scarification and monitored post infection. (A) Survival over time was observed and the significance of the percentage survival between female and male mice were analyzed using Kaplan-Meier survival analysis. (B) Clinicals scores were calculated by scoring lesion, neurological, and eye phenotypes as outlined in table 1. (C) Percent weight compared to pre-infection weights. Student T test (B-C). *p<0.05. The mean and SEM are represented. N= 13 females, 8 males.

After at least 28 days post-infection, ganglia were explanted and incubated in the presence of the PI3-kinase inhibitor, LY294002, with and without ACV. Values from both male and female mice were combined (Fig. 3A-D). Quantification of viral DNA loads showed that the copy number of viral genomes stayed constant up to 15 hours post-stimuli and were not affected by the presence of ACV, indicating that detectible viral genome synthesis did not occur in this time period. In contrast, a robust induction of viral lytic gene expression occurred by 5h post-stimuli, as inducted by an increase in IE (ICP27), E (ICP8) and late (gC) mRNA copy number. For all gene classes examined, the increase in copy number was 100-fold for IE mRNA (Fig. 3B), 20-fold for E mRNA (Fig. 3C), and 10-fold for L mRNA (Fig. 3D) 5 hours post explant. The robust increase in late gene expression in the absence of detectible viral genome synthesis suggested that late gene expression could occur even prior to DNA replication. To further support this conclusion, late gene induction occurred to equivalent levels even in the presence of ACV. The inclusion of ACV did prevent genome synthesis at 20h post-stimuli (Fig. 1A), indicating that the ACV was capable of acting on the explanted ganglia. Therefore, these data indicate that the induction of lytic gene expression following PI3-kinase inhibition in explanted sensory neurons resembles at least one feature of Phase I gene expression as viral late gene expression occurred independently of viral DNA replication.

**Figure 3:**
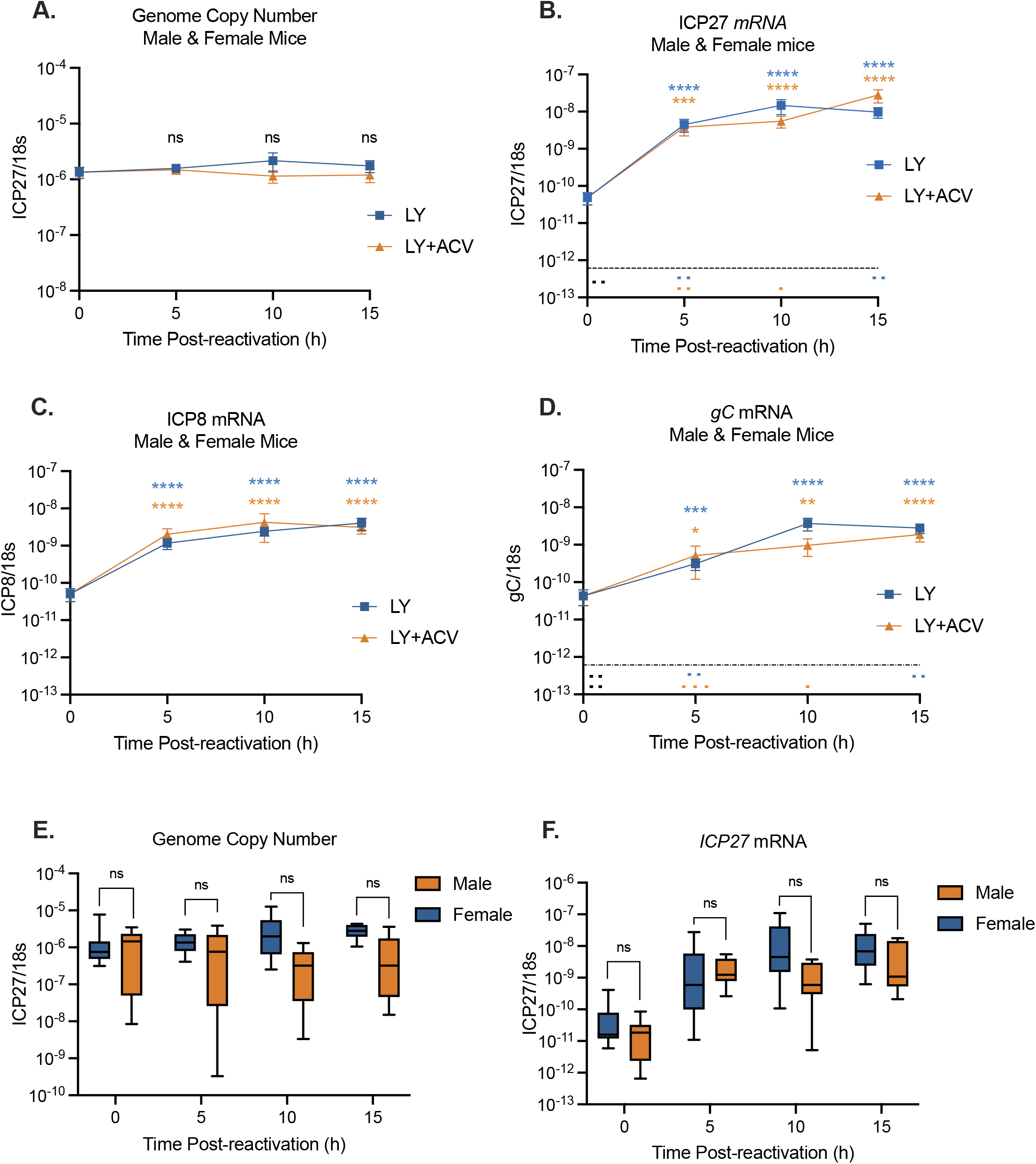
PI3K inhibition of explanted trigeminal ganglia induces rapid lytic gene expression in the absence of detectible genome synthesis Latently infected TGs from male and female mice were reactivated for 5,10 or 15 hours with LY294002 in the presence and absence of acyclovir. (A) Viral genome copy number was quantified by PCR. Viral gene expression also was quantified by RT-qPCR for (B) immediate early (*ICP27*), (C) early (*ICP8*), and (D) late (*gC*) genes. The average genome copy number (E) and ICP27 mRNA transcripts (F) for male and female mice were calculated. Transcript copy number was normalized to cellular control (18s rRNA). Limit of detection indicated by black dashed line. N≥19 biological replicates (A-D). Mann-Whitney U (**A-F**). *p<0.05, **p<0.01, ***p<0.001. ****p<0.0001. The means and SEMs are represented.

To confirm that reactivation had no sex-dependent effects we performed quantification of viral lytic gene expression in both male and female mice. We found no difference in viral genome copy number (Fig. 3E) or ICP27 lytic gene expression (Fig. 3F) at any time point. We therefore concluded that the observed phenotypes in lytic gene expression were sex-independent.

### PI3-kinase Inhibition in Combination with Axotomy Triggers Rapid Lytic Gene Expression in Sympathetic Neurons *Ex Vivo*

Previous studies investigating Phase I gene expression *in vitro* have largely used sympathetic neurons, where the induction of lytic gene expression has been found to occur 15-20h post stimuli (25-27). However, we found that lytic gene expression induced ex vivo from sensory neurons was robustly induced by 5h post-stimulus (Fig 2). Therefore, to determine whether the enhanced kinetics of reactivation observed *ex vivo* from sensory neurons in the trigeminal ganglia could result from the use of different neuronal types, we investigated reactivation *ex vivo* from the superior cervical ganglia (SCGs). Latently infected SCGs were explanted and incubated in the presence of the PI3-kinase inhibitor, LY294002. Quantification of viral DNA loads revealed that the copy number of viral genomes remained constant at 5 hours after stimulation, indicating that no detectable viral genome synthesis occurred during this time (Fig. 4A). In concordance with our sensory ganglia data (Fig. 1), there was a significant 10-fold increase in viral genome copy number at 20 hours post excision (Fig. 4A). By 5h post-stimuli a robust stimulation of viral lytic gene expression had occurred, as evidenced by a 10-fold increase in IE (*ICP27*, Fig 4B) and a 20-fold increase in late (*gC*) mRNA copy numbers (Fig. 4C). This indicates that the increased kinetics observed *ex vivo* in sensory neurons was not due to different neuronal subtypes and instead likely results from either the combination of triggers or as a result of *in vivo* infection.

**Figure 4:**
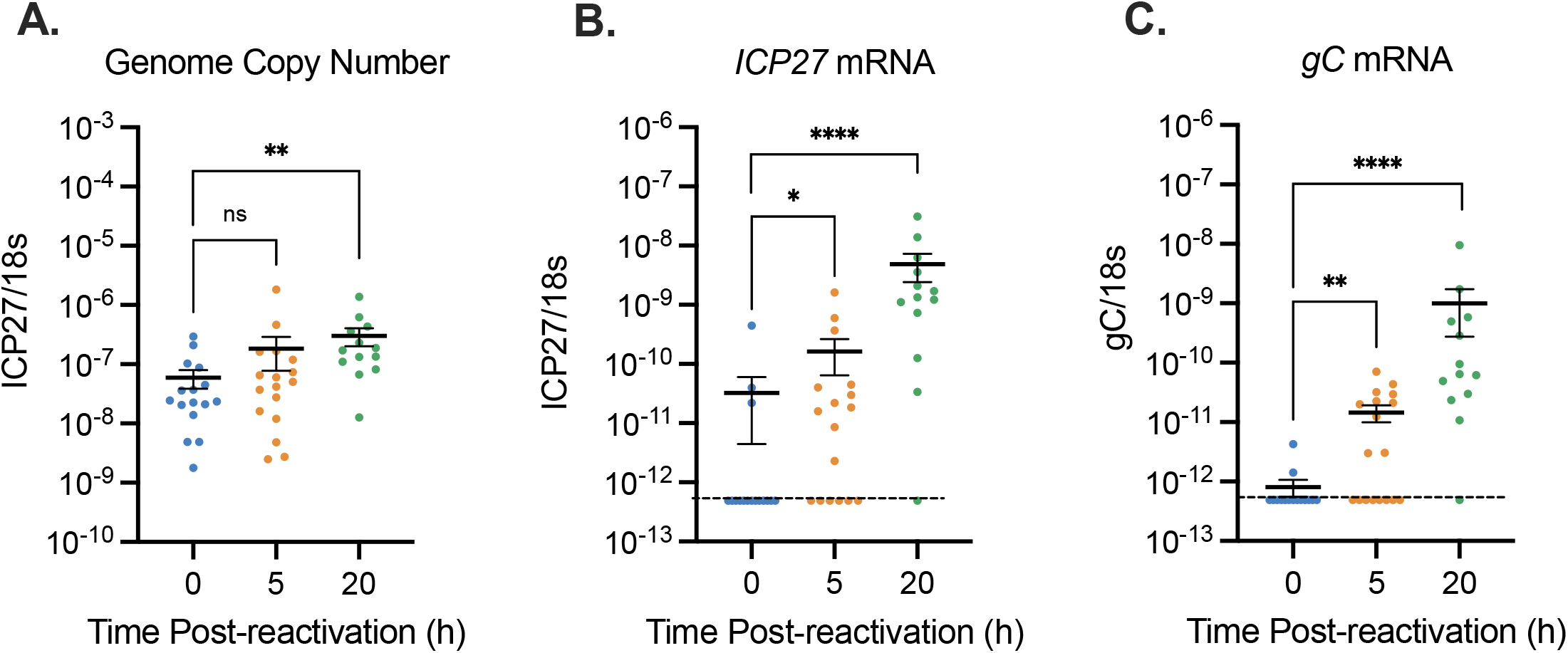
PI3K inhibition of explanted superior cervical ganglia induces rapid lytic gene expression in the absence of detectible genome synthesis. Latently infected SCGs from female mice were reactivated for 5 or 20 hours with LY294002. (A) Viral genome copy number was quantified by PCR. Viral gene expression was quantified by RT-qPCR for (B) immediate early (*ICP27*), and (C) late (*gC*) genes. Transcript copy number was normalized to cellular control (18s rRNA). Limit of detection indicated by black dashed line. N≥12 biological replicates (A-C). Mann-Whitney U (**A-C**). *p<0.05, **p<0.01, ***p<0.001. ****p<0.0001. The means and SEMs are represented.

### Analysis of preformed virus production in sensory neurons *ex vivo*

Based on our data that viral late gene expression was detectible by 5 hours and robustly expressed by 10 hours post-explant, we investigated when preformed infectious virus could be detected in the explanted ganglia. The rationale for this was that Phase I lytic gene expression occurs in cultured neurons before the production of *de novo* virus (25). To ensure that we detected only preformed virus and not virus produced from remaining intact cells, the ganglia were homogenized and subjected to three rounds of sonication and two cycles of freezing and thawing. Using a viral stock of known titer, we confirmed that that this procedure did not result in a detectible loss in viral titer (data not shown). Infectious virus was robustly detected at 20h post-explant in seven out of the eight ganglia tested (Fig. 5). At 15 hours, virus was only detected in four out of eight ganglia tested and at 10 hours only two ganglia had detectible virus. No infectious virus was detected at 7 hours post-explant. These findings suggest that a detectable increase in late gene expression occurs 10 to 15 hours before infectious virus production, observed at 20 hours, in a population of reactivating ganglia.

**Figure 5:**
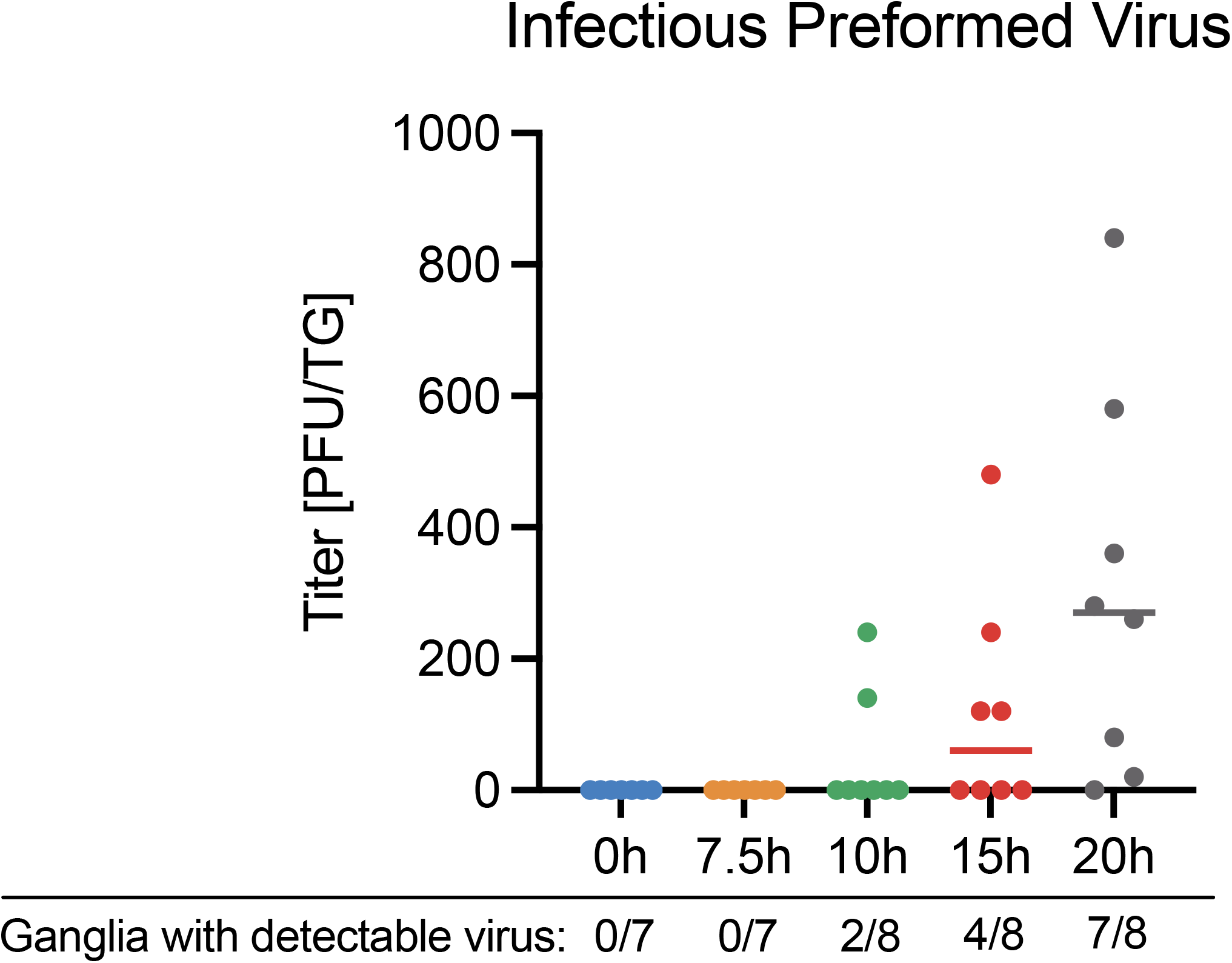
Robust detection of pre-formed virus occurs 20 hours post excision Latently infected TGs from female mice were reactivated for 0, 7, 10, 15, or 20 hours with LY294002 and titers of pre-formed virus quantified on Vero cells. Number of ganglia with detectible virus is displayed beneath the X axis. N≥7 biological replicates. The median titer is represented.

### DLK activity is required for reactivation *ex vivo*

Previously, we have found that the neuronal cell stress protein dual leucine zipper kinase (DLK) is required for HSV reactivation and acts to induce Phase I lytic gene expression. To determine whether DLK was also required for induction of lytic gene expression in sensory neurons reactivated *ex vivo* by PI3K-inhibition/axotomy, the DLK inhibitor GNE-3551 was added to the explanted ganglia and viral RNA was extracted at 5h post-reactivation. Inclusion of the DLK-inhibitor resulted in *ICP27, ICP8*, and *gC* mRNA (Fig. 6 A-C) levels that were equivalent to the un-reactivated samples and significantly decreased compared to the reactivated ganglia that were not treated with the DLK-inhibitor. These data indicate that DLK is required for reactivation from sensory neurons induced by the combined trigger of explant and PI3-kinase inhibition.

**Figure 6:**
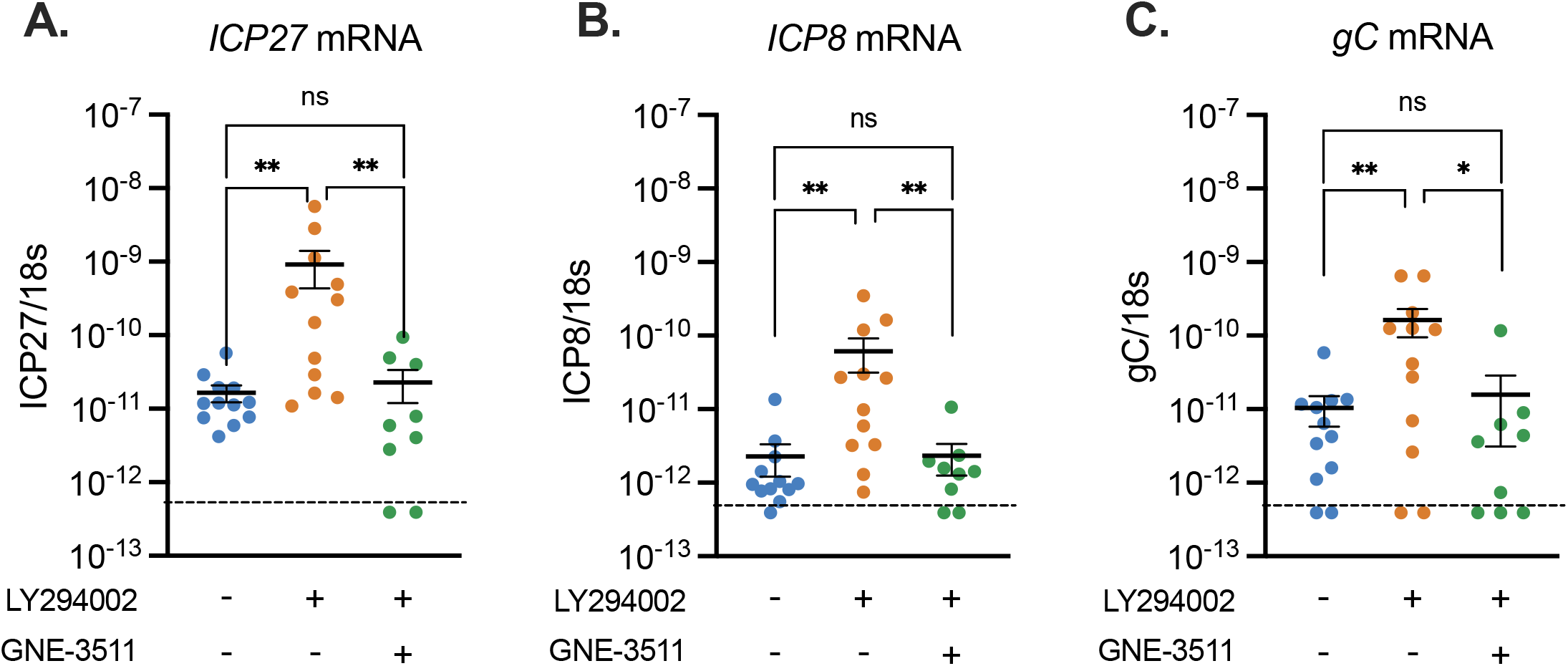
DLK activity is required for the induction of lytic gene expression following explant/PI3-kinase inhibition of latently infected TG. Latently infected TG were explanted and incubated with LY294002 and the DLK inhibitor GNE-3511 (8 μM) for 5 hours post excision. Viral gene expression was quantified by RT-qPCR for immediate early (*ICP27*) (A), early (*ICP8*) (B), and late (*gC*) (C) genes. Limit of detection indicated by dashed line. n=12 biological replicates. Mann-Whitney U test. *p<0.05, **p<0.01. Individual biological replicates along with the means and SEMs are represented.

### Lytic gene induction upon axotomy along with PI3-kinase inhibition is independent of histone H3 histone lysine 9 and lysine 27 demethylase inhibitors

HSV promoters are known to be enriched with histone H3 di- and trimethyl at lysine 9 (H3K9me2/3) and histone H3 trimethyl at lysine 27 (H3K27me3) during latency. The removal of restrictive histone modifications, specifically the aforementioned methylation marks, was previously shown to be essential for full reactivation of HSV *in vitro*, yet was not required for initial lytic gene expression during Phase I of reactivation (27, 30). To determine if initial lytic gene expression was independent of histone demethylation *ex vivo*, we used OG-L002 (60 μM), a drug that inhibits the histone lysine 9 demethylase LSD1, and GSK-J4 (20 μM), a molecule that inhibits the histone lysine 27 demethylases UTX and JMJD3. HSV reactivation has been demonstrated to be inhibited by both of these inhibitors (27, 30, 44, 45). Five hours post excision, during Phase I of reactivation, the addition of OG-L002 and GSK-J4 had no effect on the induction of ICP27, ICP8, or gC mRNA (Fig. 7A-C). These results indicate that histone demethylase activity is not required for the initial induction of lytic gene expression induced by axotomy and PI3-kinase inhibition *ex vivo*.

**Figure 7:**
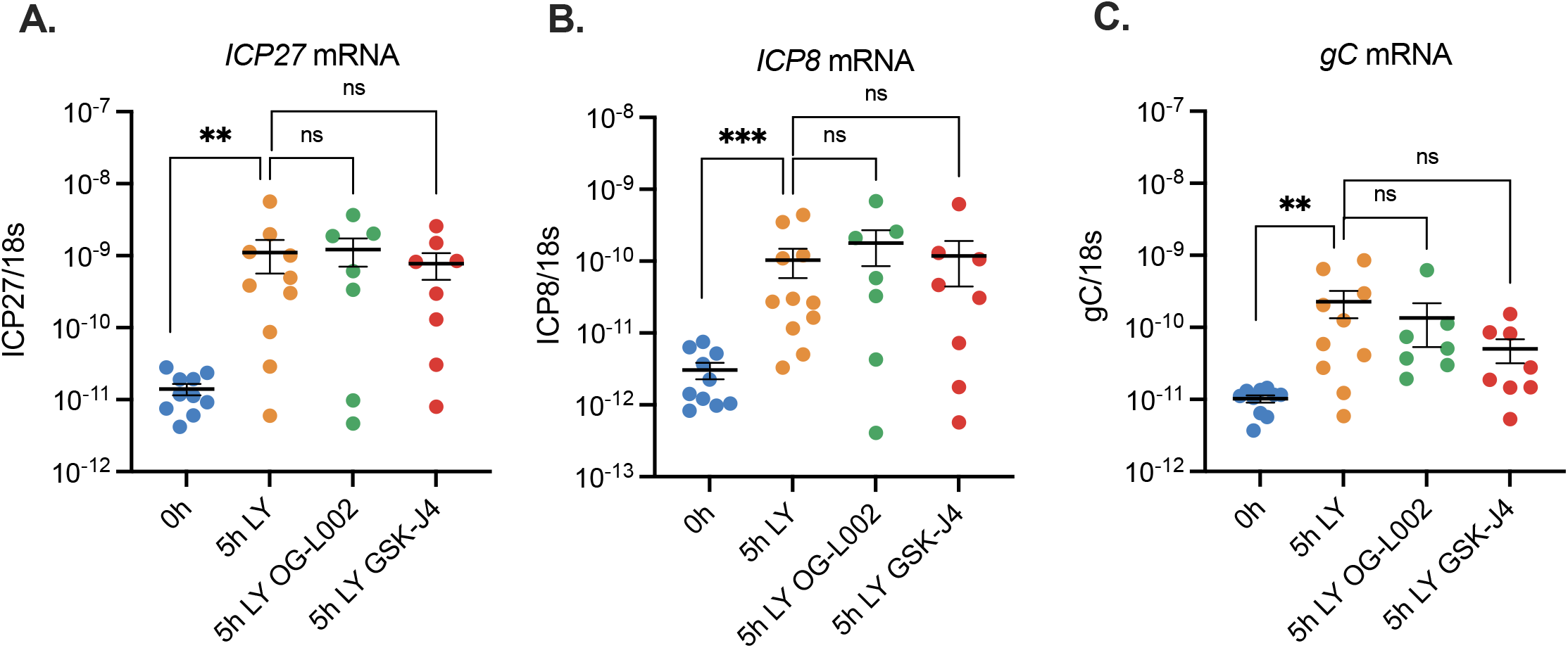
Lytic gene induction following axotomy/PI3-kinase inhibition is unaffected by histone demethylase inhibitors. Latently infected TG were explanted and incubated with LY294002 along with the LSD1 inhibitor OG-L1002 (50 μM) or GSK-J4 (20 μM). Viral gene expression was quantified by RT-qPCR for immediate early (*ICP27*) (A), early (*ICP8*) (B), and late (*gC*) (C) genes. Mann-Whitney U Test, **p<0.01, ***p<0.001. Individual biological replicates along with the means and SEMs are represented.

Together, our results demonstrate that features of Phase I gene expression, namely late gene expression in the absence of viral DNA replication and lytic gene expression in the presence of histone demethylase inhibitors, can occur from ganglia isolated from infected mice. In addition, *ex vivo* reactivation was dependent on DLK activity.

## Discussion

*In vitro* models of HSV latency are incredibly powerful for the study of molecular mechanisms HSV latency and reactivation as well as the contribution of host factors and viral factors during specific stages of infection. *In vitro* models can be used to investigate how the host immune response can modulate latent infection and reactivation. Using these models, unique aspects regarding entry into lytic gene expression during reactivation have been uncovered, including the dependence on DLK/JNK for reactivation, the ability of late gene expression to occur in the absence of DNA replication, and transcription despite the presence of histone demethylase inhibitors. Therefore, it was highly important to validate these patterns of gene expression from ganglia infected *in vivo*. Here we have confirmed that the features of gene expression observed during Phase I reactivation *in vitro* also occur following reactivation *ex vivo* when PI3k-inhibition and axotomy is used as a trigger.

There are certain caveats to our study, especially relating to the ability of inhibitors to act on intact ganglia. However, we show that the DLK inhibitor, PI3-kinase inhibitor and acyclovir can all act on intact ganglia. Therefore, we think it unlikely that the histone demethylase inhibitors used were unable to penetrate the tissue and have an effect on viral lytic gene induction. The concentrations of histone demethylase inhibitors were higher than what we have previously observed to inhibit full HSV reactivation in primary neuronal models (27, 30). We have shown that OG-L002 can inhibit forskolin mediated reactivation at 30 μM, here we used 60 μM. Similarly, we have shown that GSK-J4 can inhibit both forskolin and LY294002 induced reactivation at 3 μM; here we used it at 20 μM. Therefore, we think it unlikely that the concentrations used were too low to inhibit the histone demethylases. We did try higher concentrations that resulted in loss of viral genomes at 5h, indicating potential neuronal toxicity at these higher concentrations.

Our study did result in the discovery of a unique phenotype of Phase I of reactivation when observed *ex vivo*. Unlike our previous *in vitro* studies, we found that Phase I of reactivation that occurred *ex vivo* had enhanced kinetics of lytic gene induction upon axotomy and PI3-kinase stimulation. We observed this enhanced phenotype in both sensory and sympathetic ganglia. A paralleled kinetic phenotype in both neuronal subtypes was validated in a novel model of latency and reactivation utilizing a virus defective in cell spread, in which our lab discovered that reactivation kinetics *in vitro* were equivalent in sensory and sympathetic neurons (Dochnal et al, manuscript in preparation). Therefore, we posit that the most likely explanation of the difference in reactivation kinetics between *in vitro* and *ex vivo* studies is the combined trigger of axotomy and PI3-kinase inhibition. It has previously been demonstrated that when nerve growth factor deprivation is paired with axotomy, treated neurons undergo cell death more quickly than untreated neurons. (46). In addition, both PI3-kinase inhibition and axotomy result in activation of DLK (32, 47). However, there are some differences in how DLK is activated and the resulting downstream response in response to the two stimuli. Following axotomy, DLK is rapidly activated at the proximal segment of the severed axon in response to loss of cytoplasmic integrity and a calcium influx results from cytoplasmic membrane rupture to ultimately result in transcription of genes involved in axon regeneration (48). The exact mechanisms of DLK activation upon loss of NGF signaling that mediates neuronal cell death or axon pruning is not fully understood. It is known that in response to NGF deprivation, DLK protein levels are stabilized, DLK is phosphorylated (49), and this phosphorylation results in the downstream activation of JNK, c-Jun, and other transcription factors. The activation of these transcription factors promotes the expression of pro-apoptotic genes (50), although in mature neurons there are multiple brakes downstream that prevent apoptosis (51). It is therefore conceivable that these two independent pathways to DLK activation may converge follow the dual trigger of axotomy/PI3-kinase inhibition to result in more rapid and robust DLK activation and HSV reactivation.

Consistent with cellular pathways converging the enhance DLK activity, we observed a robust induction of lytic gene expression at 5h post reactivation. For the IE gene ICP27 this was an approximate 100-fold induction, compared to the 5-20-fold increase often observed in an *in vitro* mouse model with PI3-kinase inhibition (27). The use of an *ex vivo* model system will therefore be incredibly powerful for studying the mechanisms of gene expression that occur during Phase I. We have previously observed a histone methyl/phospho switch on lytic promoters during Phase I reactivation (27), which permits lytic gene expression independently of recruitment of histone demethylase enzymes. There is evidence that the methyl/phospho switch permits gene expression because repressive histone readers (for example HP1) are no longer capable of interacting with the methylated residue on histone H3K9 because phosphorylation at Serine 10 occludes binding (52, 53). Because of the robust lytic gene induction *ex vivo*, and because we have now shown that the gene expression both *in vitro* and *ex vivo* is DLK-dependent and histone demethylase independent, the complementary models can now be used answer key questions on the mechanism of viral lytic gene induction during Phase I reactivation.

## Acknowledgements

We thank David Knipe (Harvard Medical School) for the KOS virus. This work was supported by R01NS105630 (ARC), The Owens Family Foundation (ARC), NIH/NIAID T32AI007046 (ALW, JBS), NIH/NEI F30EY030397 (JBS) and NIH/NIGMS T32GM008136 (SAD).

